# You’d better decide first: overt and covert decisions improve metacognitive accuracy

**DOI:** 10.1101/470146

**Authors:** Marta Siedlecka, Zuzanna Skóra, Borysław Paulewicz, Michał Wierzchoń

## Abstract

How can we assess the accuracy of our decisions? Recent theoretical and empirical work suggest that confidence in one’s decision is influenced by the characteristics of motor response in a preceding decisional task. In this paper we present experiment designed to test whether decision itself can also inform confidence and therefore increase its accuracy. We tested 143 participants who solved an anagram-solving task in one of 3 conditions: participants either rated their confidence immediately after responding to the anagram task (overt decision), rated their confidence immediately after making a decision but without overt response (covered decision), or rated their confidence before both deciding and responding. The results showed significant relationship between decision accuracy and confidence level in each condition, however this relation was stronger when confidence rating followed decision, either covert or overt. We argue that completing a decisionmaking process increases metacognitive accuracy.

---

How can we assess the accuracy of our decisions? This issue has been studied for a long time, but it is still a subject of intense debate (Fleming & Daw, 2018; Gajdos, Fleming, Saez Garcia, Weindel, & Davranche, 2018; Samaha, Iemi, & Postle, 2017; Siedlecka, Skóra, Fijałkowska, Paulewicz, Timmermans, & Wierzchoń, 2018; Wokke, Cleeremans, & Ridderinkhof, 2017). For the last decades this debate was focused on the issue of whether confidence in one’s decision is based on the same information as decision itself, such as the quality of stimulus or memory trace (e.g. Galvin, Podd, Drga, & Whitmore, 2003; Higham, Perfect & Bruno, 2009; Vickers & Lee, 1998) or whether it is also informed by additional, post-decisional evidence (Graziano, Parra & Sigman, 2015; Hilgenstock, Weiss & Witte, 2014; Moran, Teodorescu, & Usheret, 2015; Pleskac & Busemeyer, 2010; Ploran, Nelson, Velanova, Donaldson, Petersen, & Wheeler, 2007). In recent years the focus of the discussion has shifted towards post-decisional processes, resulting in growing theoretical (Fleming & Daw, 2018) and empirical interest in the ways action can influence confidence. Numbers of studies have shown that confidence level is affected by the characteristics of motor response associated with decision-making (Faivre, Filevich, Solovey, Kühn, & Blanke, 2018; Fleming, Maniscalo, Ko, Amendi, Ro & Lau, 2015; Gajdos et al., 2018; Kiani, Corthell & Shalden, 2014). For example, it has been shown that confidence is influenced by motor-related neural activity that is associated with response that is being prepared. Fleming and colleagues (2015) applied transcranial magnetic stimulation (TMS) to the premotor cortex area associated with either a chosen or unchosen response and observed that participants reported lower confidence in their perceptual decision when the stimulation was incongruent with their actual response. Another study has shown that participants were more confident in responses preceded with stronger motor preparation (measured with electromyograph), no matter whether they were congruent or incongruent with the actual response (Gajdos et al., 2018, see also Siedlecka, Hobot, Skóra, Paulewicz, Timmermans, & Wierzchoń, 2018). Once response is given confidence might be informed by the results of error-monitoring (Bold & Yeung, 2015; Scheffers & Coles, 2000). For example, Boldt & Yeung (2015) observed that error-related brain activity that is present after an erroneous task response is associated with lower confidence in the preceding perceptual decision.

The discovered relationships between response to a perceptual stimuli and the level of confidence might not be surprising when taking into consideration data from neurophysiological studies suggesting that in decision tasks, where stimuli characteristics are associated with a specific action, making a decision might be indistinguishable from action selection and preparation (Gold & Shadlen, 2003; Hernández, Zainos, & Romo, 2002; Wyss, König, & Verschure, 2004). However, studies also reveal the possibility of a general decision-making system, that integrates evidence from different sources until a decision can be made (Heekeren, Marrett, Bandettini, & Ungerleider, 2004; Heekeren, Marrett, & Ungerleider, 2005; Heekeren, Marrett, Ruff, Bandettini, & Ungerleider, 2006). This system, thought to be located in the left posterior dorsolateral prefrontal cortex, is engaged in decision-making process irrespectively of the type of stimuli and response modality.

The process of decision-making has not received much attention in the context of confidence research. Although making a decision and carrying out motor response might provide different type of information for confidence, it is hard to separate them either experimentally or conceptually. For example, although reaction time in a decisional task is one of the most commonly measured correlates of confidence (Dougherty, Scheck, Nelson, & Narens, 2005; Kelley & Lindsay, 1993; Mealor & Dienes, 2013; Petrusic & Baranski, 2003), it is not clear whether it provides information about the stimulus difficulty (Pleskac & Busemeyer, 2010; Van Zandt and Maldonado-Molina, 2004; Vickers & Lee, 1998), decision-making process (Kiani et al., 2015) or motor response preparation and execution. Similarly, when measuring task performance researchers usually do not differentiate between decision and motor response accuracy, although participants might be aware that they carried out different response to the one they intended (e.g. when they report committing an error, Boldt & Yeung, 2015; Scheffers & Coles, 2000).

In this study we experimentally separated decision from its motor execution to test the effect of decision itself on metacognitive accuracy, that is to see whether it influences the extent to which a person’s confidence predicts their decision correctness. We used an anagram task, in which response is not directly linked with the physical characteristics of a stimulus as we assumed it involves more higher cognitive processing that simple perceptual decisions. In order to separate decision from its motor execution we introduced an overt and covert decision conditions. In the Overt decision condition participants immediately expressed their decision with motor response and then rated confidence in that response. In the Covert decision condition participants were asked to decide before rating their confidence but they carried out motor response afterwards. In the No-decision condition participants were first asked to rate their confidence and then to respond to the anagram task. We compared metacognitive accuracy between all three conditions. We expected that metacognitive accuracy would be higher when confidence ratings follow Covert decision (decision without motor response) compared to the No-decision condition, which would indicate that the results of decision-making process provide crucial information for confidence judgments we should expect. If however decisions have to be carried out to affect metacognitive accuracy, this parameter should be highest in the Overt decision condition than in two other conditions.

## Methods

### Participants

One hundred and forty-three participants^1^, 30 male, aged 19-28 (*M*=21.31, *SD*= 1.78) took part in the experiment in return for a small payment (about 15 PLN). All participants had normal or corrected to normal vision and gave written consent to participation in the study. The ethical committee of the Institute of Psychology of the Jagiellonian University approved the experimental protocol.

### Materials

The experiment was run on PC computers using E-Prime. For the purpose of the study, 80 Polish nouns, containing 7-10 letters were chosen. The words were paired so that 30 pairs differed by just 1 letter (in English this could be: SENATOR-TOASTER), 9 pairs differed by 2 letters (e.g., RESTAURANT-TRANSLATOR), and one pair that differed in 3 letters, which could be either exchanged or added. The anagrams were made by randomly mixing the letters of one word in a pair. Three judges chose one letter string for each anagram that was least similar to any word and did not contain any syllables included in the solution or target word. The list of anagrams to solve was the same for all participants but different solutions (i.e., correct or incorrect) were suggested. We used a confidence scale to measure confidence of participants’ decision (“How confident are you that you will make the right decision?” or “How confident are you that you made the right decision?”). The answers were: “I am guessing,” “I am not confident,” “I am quite confident,” and “I am very confident.”

### Procedure

Participants were tested in small groups in a computer laboratory and randomly assigned to one of three groups. The outline of the procedure is presented in Figure 1. Each trial started with a fixation-cross appearing for 1s and followed by an anagram written in capital letters. Each anagram was presented for 20s. Then it was masked by $$$$$$$$$$ symbol for 200 ms and followed by a 200 ms blank screen. A target word was presented for 350 ms (so as participants could not easily compare or “rearrange” the letters). Then, in Overt decision condition a question “solution?” was presented in the centre with two options “yes” and “no” on both sides. After making the decision participants were asked to rate their certainty on the confidence scale. In the No-decision condition participants were first asked to rate their confidence and then to respond whether a target word was an anagram solution. In the Covert decision condition participants were first presented with a question “decision?” and asked to signal when they have made their decision. Afterwards they rated their confidence and then responded “yes” or “no” to the “solution?” question. Participants in this group were asked not to change their minds and just to report decision they made before confidence rating^2^. Participants in both Overt decision and No-decision groups were given up to 3000 ms for response and for confidence rating. In Covert decision time available for covert decision and confidence rating was 3000 ms and for the following response it was limited to 1500 ms. In all groups participants could take a break after each trial and proceed to next trial by pressing the Space key.

**Figure 1:**
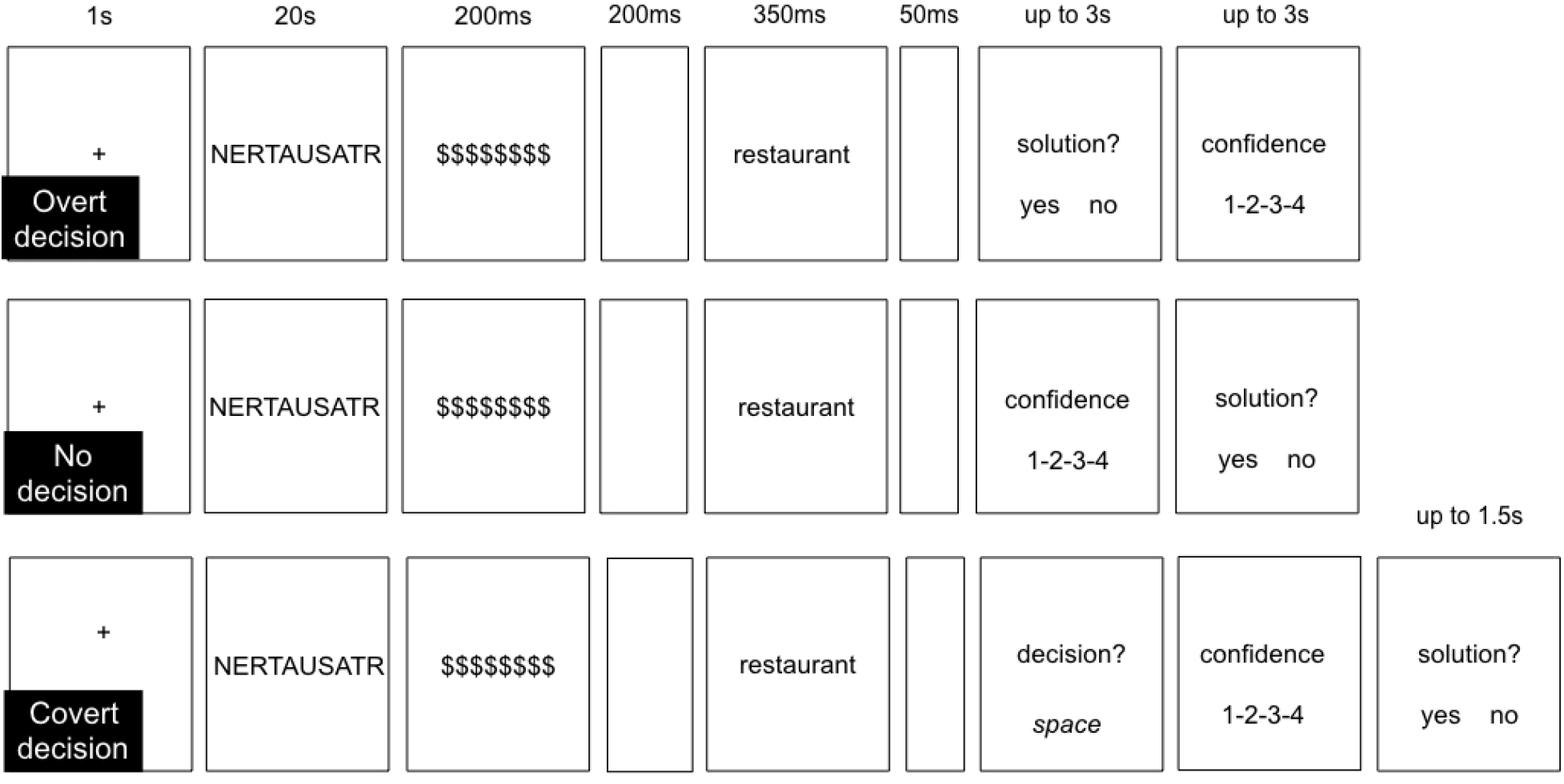
The three conditions of the anagram task. Overt decision: participants first respond in the anagram task and then rate their confidence; No decision: participants first rate their confidence and then respond in the anagram task; Covert decision: participants first make anagram-related decision, then they rate their confidence and finally they carry out anagram-related response.

Participants responded “yes” and “no” by pressing “o” or “p” keys with their right index and middle fingers. For half of the participants “yes” was presented on the left side of the screen and corresponded to the left key (“o”) and “no” was presented on the right side of the screen and corresponded to the right key (“p”), and for half of the participants the arrangement was the opposite. The keys corresponding to the “yes” and “no” response were covered with a green and red stickers, respectively. The confidence level was reported with keys (“1,” “2,” “3,” and “4” using the left hand (from “1” representing “I am guessing” to “4” representing “I am very confident”). Covert decision was signalled by pressing space bar with a thumb of the right hand.

The experiment started with two training trials during which participants were shown two examples of simple anagrams. There were 40 experimental trials during which anagrams were presented in random order. For each participant half of the target words were the correct solutions.

## Results

Prior to data analysis we removed all non-complete trials, that is trials in which at least one reaction was missing (anagram-related response, confidence rating or covert decision, in total 12% of all trials) and trials in which reaction times were lower than 150 ms (based on discontinuity in reaction time distribution). Next we fitted a Bayesian mixture model to the number of complete trials (see e.g. Lee & Wagenmakers, 2014). We assumed that each participant comes from one of two distributions that differ in the probability of any given trial being complete. Thirteen participants, who completed less than 67,5% of trials were omitted from further analyses (it was a posteriori more likely that they belong to the distribution with 50% probability of not completing a trial). There were 47 participants left in Overt decision group, 38 participants in Covert decision group and 45 participants in No decision group.

The accuracy in anagram task was above chance level. An average accuracy level in each group equalled: Over decision — 79%, Covert decision — 76%, No decision — 75%. We observed no statistically significant differences in accuracy between groups (|z| < 1.76, *p* > .08), neither found we significant differences in d’ (|z| < 1.84, *p* > .07) or bias (|z| < 1.38, *p*> .17).

Metacognitive accuracy was operationalized as the extent to which confidence ratings predict response accuracy in the anagram task (Norman & Price, 2015). The mixed logistic regression models were fitted using the lme4 package in the R Statistical Environment (Bates, Maechler, Bolker, & Walker, 2015; R Core Team, 2015) using standard (0/1) contrast coding. In order to test the effect of condition on metacognitive accuracy we fitted a model that included fixed effects of confidence rating (4 levels), condition (3 levels) and their interactions, as well as random effects of participant-specific intercept and slope. Confidence ratings were centred on the lowest value (“guessing”) and the basic condition was Overt decision (task response followed by confidence rating). Therefore, the regression slope reflects the relation between accuracy and confidence rating (that is metacognitive accuracy) while the intercept informs about performance level when participants report guessing. Statistical significance was assessed with the Wald test.

The results are presented in Table 1 and on Figure 2. The first row (intercept) shows an estimate of the average accuracy (on the logit scale) in the baseline condition (Overt decision, lowest rating = “guessing”). When reporting guessing participants in Overt decision condition did not perform significantly different from chance level. The second and third row shows that performance of participants in Covert decision and No-decision conditions, when they reported guessing, did not differ significantly from performance in Overt decision (for “guessing”). The fourth row estimates regression slope in the Overt decision group and shows that the relationship between Accuracy and Confidence in this condition is statistically significant. The fifth row shows that no significant difference was found between the slopes in Covert decision and Overt decision. The last row indicates that the slope is significantly lower in No-decision condition compared to Overt decision. After reparametrisation of the model (No-decision as baseline condition) we found no significant difference in the intercept between No-decision and Covert decision conditions (*z* = -1.54, *p* = .12). However, according to the one-tailed test, the regression slope is lower in No-decision condition than in Covert decision (*z* = 1.93, *p* = .025).

**Table 1.** Regression coefficients for the logistic regression mixed model for accuracy

| **Number of participants: 130**<br>**Number of observations: 4532** | **Estimate** | **SE** | **z** | **p** |
| --- | --- | --- | --- | --- |
| Intercept | -0.08 | 0.14 | -0.57 | .56 |
| Condition: Covert decision | -0.19 | 0.22 | -0.84 | .40 |
| Condition: No-decision | 0.15 | 0.20 | 0.77 | .44 |
| Confidence | 0.8 | 0.08 | 10.28 | < .001 *** |
| Confidence: Covert decision condition | -0.03 | 0.12 | -0.24 | .81 |
| Confidence: No-decision condition | -0.25 | 0.12 | -2.33 | .02 * |
| Likelihood ratio: $\chi^2(7) = 340, p < .001$ | | | | |
| *$p < .05$; **$p < .01$; ***$p < .001$ | | | | |

**Figure 2:**
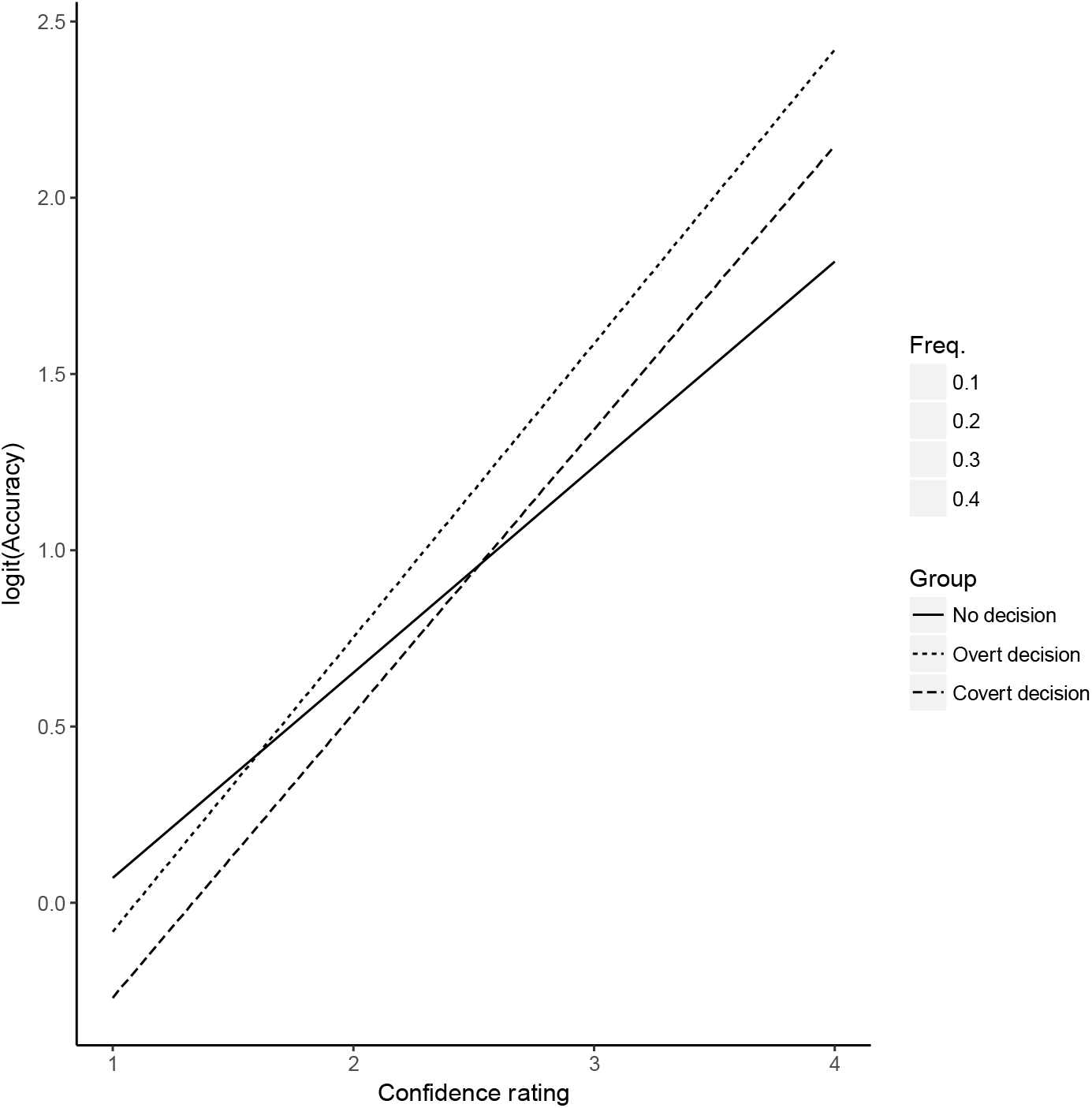
Model fit for the relationship between accuracy and confidence ratings in each condition. The size of filled circles represents frequency of each confidence rating.

To test whether Overt decision and Covert decision conditions are similar in terms of the relationship between accuracy and confidence ratings, we compared the fit of the model with a second model, in which those two conditions are merged (Covert decision and Overt decision). We found no significant differences between models’ fit (*χ*2(2) = 2.58, *p* = 0.27). According to the second model participants in neither groups differed in intercept (*z* = 1.27, *p* = .2). However, regression slope was significantly lower in No-decision condition than in the other conditions (*z* = -2.47, *p* = .01).

Additionally, using mixed logistic regression model, we tested whether the latency of covert decision was related to the subsequent accuracy of anagram-related motor response (using mixed logistic regression). As expected, covert decision took longer when the subsequent response was incorrect (t(37) = -2.46, *p* = .02), which suggests that the covert decision time reflects underlying decisional process. Similarly, response time was longer for incorrect responses in all conditions (Overt decision: z = -5.40, *p* < .001; No-decision: *z* = -4.02, *p* < .001; Covert decision: *z* = -4.44, *p* < 0.01). Using linear mixed model we also estimated the relationship between response time and confidence ratings. The results showed that the later the response was given the lower confidence level was in each condition (Overt decision: *t*(79) = -11.8, *p* < .001; No-decision: *t*(111) = -8.76, *p* < .001; Covert decision: *t*(331) = -6.44, *p* < .001). Similarly, longer Covert decision time was associated with lower confidence: *t*(31) = -5.8, *p* < .001.

We carried out additional analyses to find out whether the difference between conditions in metacognitive accuracy could have come simply from the time of confidence rating (measured from target word presentation). We created a new variable, Confidence rating time, which in the No-decision condition equalled confidence rating RT, in the Overt decision equalled the sum of response RT and confidence rating RT, and in the Covert decision condition equalled the sum of covert decision RT and confidence rating RT. Since the conditions differed in confidence rating time (for procedural reasons) this analyses were carried within conditions. We found a significant relation between Confidence rating time and metacognitive accuracy only in No-decision condition (*z* = -4.2, *p* < .001): the later the rating was given the less accurate it was. We did not observe this relation in any of the other conditions (*p* > .16).

## Discussion

In this experiment we separated decision from its motor execution, to find out whether decision alone provides information that can increase metacognitive accuracy. We compared metacognitive accuracy between the three conditions: Overt decision, in which participants rated their confidence immediately after responding to the target word, Covert decision, in which participants rated their confidence after decision but before motor response, and No-decision, in which participants rated their confidence before responding to the target word. The results of the experiment showed that although the level of confidence correlated with the level of performance in all three conditions, metacognitive accuracy was higher in the conditions in which decision preceeded confidence rating compared to the No-decision condition. We did not find statistically significant difference in metacognitive accuracy between Covert and Overt decision conditions. The effect of lower metacognitive accuracy in the No-decision condition could not be explained by the smaller amount of time available for confidence judgment in this condition. The relation between rating time and metacognitive accuracy was negative: the earlier confidence ratings were given the more accurate they were.

The result of this experiment, showing that metacognitive accuracy was higher in the Covert decision condition compared to No-decision, suggests that decision improves confidence accuracy even when it is not expressed with decision-specific motor response. In our previous studies with anagram task (Siedlecka, Paulewicz, & Wierzchoń, 2016), memory task (Siedlecka et al., 2018) and perceptual decision task (Wierzchoń, Paulewicz, Asanowicz, Timmermans, & Cleeremans, 2014) we observed higher metacognitive accuracy when confidence judgments or visibility judgments followed task response compared to the conditions in which they preceded that response. However, in none of those studies decision was separated from motor response. Moreover, although there is data showing that action-related characteristics influence or correlate with the level of confidence, there has been almost no empirical evidence suggesting that motor-related information affects metacognitive accuracy (but see Fleming et al., 2015).

The result showing that Covert condition increased metacognitive accuracy compared to the No-decision condition does not support the so-called single-stage models of confidence, assuming that confidence and decision are based on the same information (Galvin et al., 2003; Higham et al., 2009; Vickers & Lee, 1998). What additional information is provided by the covert decision? The main difference between Covert decision and No decision conditions is that the first one requires participants to complete decisional process before assessing their confidence and thus provides information needed for this assessment. One type of such information might come from the results of comparing evidence for alternative choices available at the time of the decision (Heekeren, Marrett, & Ungerleider, 2008). Another informative aspect of a decision process is its duration. Correct responses in decision task are usually given quicker than incorrect ones (Pleskac & Busemeyer, 2010; Van Zandt and Maldonado-Molina, 2004; Vickers & Lee, 1998), and at least in some cases, response time seems to reflects decision difficulty rather than the quality of sensory evidence (Kiani et al., 2014). The results of this experiment and other studies have shown that time of response correlates with metacognitive ratings (Faivre et al., 2018; Dougherty et al., 2005; Kelley & Lindsay, 1993; Kiani et al., 2014; Koriat & Ma’ayan, 2005) and that confidence judgments are more accurate in participants whose reaction times differed most between correct and incorrect responses (Faivre et al., 2018). However, in this experiment we observed that the level of confidence was also higher for faster cover decisions, that were not expressed in a specified, decision-related response (ie. “yes” or “no”). It suggest that the time of both, decision and motor response carried information about the difficulty of matching previously presented letter string (anagram) with a target word. However only in Covert and Overt decision conditions this information was available before confidence rating. Interestingly, response time is thought to be strongly related to feelings such as ease or difficulty of processing (Dougherty et al., 2005; Koriat & Ma’ayan, 2005), which according to the experience-based view on metacognition, are highly informative for metacognitive judgements (Koriat & Levy-Sadot, 2000).

We cannot rule out the possibility that in Covert decision condition decision-related motor pogramme might have been prepared automatically but was then inhibited by a participant. In this case confidence could have been informed by motor-related information in the form of partial muscular activation, as found by Gajdos and colleagues (Gajdos et al., 2018). However, partial activation was found to increase confidence but not metacognitive accuracy. Morover, in our experiment this information would have only be available in Overt and Covert decision condition, suggesting that decisions are necessary for motor preparation, even if is is launched automatically.

Alternative explanation of the effect of decision on metacognitive accuracy comes from decision-making models based on the mathematics of quantum probability theory (Busemeyer, Wang, Townsend, 2006; Kvam, Pleskac, Yu, & Busemeyer, 2015; Yearsley & Busemeyer, 2016). According to those models, before making a decision a person’s cognitive system is not in any definite choice state but is superposed with respect to possible choices. It remains in this state until a person makes a decision and enters a definite state (Busemeyer et al., 2006; Kvam et al., 2015; Wang & Busemeyer, 2016). Quantum models are used to explain so-called order effects or interference that is observed between two consecutive decisions or judgments, where the outcome of the first one influences the other (Busemeyer et al., 2006; Moore, 2002; Yearsley & Busemeyer, 2016). For example, Kvam and colleagues (Kvam et al., 2015) observed improved confidence calibration (probability of being correct assigned to one’s response, relative to the actual proportion correct) in condition where decision-related response (“left” or “right” direction of dots movement) preceded confidence rating compared to the condition in which decision-related response was simultaneous with confidence rating (“90% confident that left”). According to the authors, making decision before confidence judgment reduces person’s uncertainty. Although in their study decision was not separated from motor response, quantum models assume a possibility of the influence of decision on confidence judgments.

Summing up, the described experiment for the first time showed that decision without a specified motor response can influence the accuracy of confidence judgment. More research is needed to establish what aspects of decision are informative for confidence and how the accuracy of decision which is not expressed with motor response is monitored by cognitive control processes.

## Footnotes

1 We aimed to test 50 participants in each experimental group. Seven participants dropped-out due to technical problems or admitting that Polish was not their native language.

2 In a pilot study we established that when participants were free to change their minds after confidence ratings, the number of reported changes was less than 5% of all trials.

